# Rugnux: from diffraction images to structure at synchrotrons and XFELs in one program

**DOI:** 10.64898/2026.10.24.761522

**Authors:** Filip Leonarski, Meitian Wang

**Affiliations:** Swiss Light Source, Center for Photon Science, Paul Scherrer Institute, Villigen PSI, Switzerland

## Abstract

Rising data rates in macromolecular crystallography outpace multi-program pipelines on computing clusters. Rugnux, a single GPU-accelerated program, turns rotation or serial still diffraction images into merged data and an initial structure. At synchrotrons, results arrive seconds after the last image; on one workstation, 207 public datasets from 36 sources took a median of 15 s each, with deposition-quality. Serial XFEL data were processed faster than the collection rate, enabling real-time structure determination.

## Main

Synchrotron macromolecular crystallography (MX) is a powerful and routine method for high-resolution structural exploration in scientific research and drug discovery^1^. Its widespread adoption relies on mature data-processing programs such as XDS^2^, HKL-2000^3^, MOSFLM^4^, and DIALS^5^, which are integral to MX beamline automation today. Serial Femtosecond Crystallography (SFX) adds time-resolved studies of structural dynamics. It also relies on automated data-processing packages, including CrystFEL^6^, and cctbx^7^.

Despite these advances in automated crystallographic analysis, the growing scale and diversity of modern MX experiments are placing increasing demands on data-processing workflows. Two developments are particularly driving this change. First, advances in synchrotron sources and hybrid-pixel detectors, including EIGER2^8^, PILATUS4^9^, and JUNGFRAU^10^, have made data collection at hundreds of frames per second routine, enabling individual synchrotron beamlines to collect hundreds of datasets per shift, and SFX to run at higher repetition rates at X-ray free electron lasers (XFELs). Second, the increasing availability of experimental structures and accurate predicted models is expanding the range of experiments that can be performed systematically. Fragment and ligand screening, the investigation of structural changes across temperatures, time points, or experimental conditions, and serial crystallography all benefit from the ability to process large numbers of datasets rapidly and determine which contain interpretable structural information. In these settings, automated data processing and, where possible, initial structure solution must keep pace with data collection to guide experiments and support timely interpretation.

Existing automated workflows, including EDNA/EDNA2^11^, autoPROC^12^, xia2^13^, fast_dp^14^ and QuickProcess2^15^, address these requirements by orchestrating established crystallographic programs within beamline pipelines. However, such workflows often depend on dedicated computing infrastructure. With high-performance computing clusters, XDS-based fast_dp can return processing results in 1-2 minutes after data collection; other data processing packages and pipelines can take substantially longer. SFX data processing optimization typically continues weeks after beamtime. Reprocessing of large modern datasets on a laboratory computer has become increasingly harder. The growing disparity between data-collection rates and accessible processing capacity therefore motivates a different approach: one that combines high throughput, autonomous decision-making and reliable crystallographic analysis without requiring a large computing cluster.

Building on the kilohertz single-image analysis in Jungfraujoch^16^ and on decades of developments in the crystallographic community, we present Rugnux, a GPU-accelerated program for crystallographic data processing designed to address these challenges. Rugnux is designed to make high-throughput rotation and still MX feasible on a single workstation rather than a dedicated computing cluster. We tested performance and quality against 207 publicly available rotation datasets from 26 synchrotron facilities and 10 laboratory sources, plus 11 serial still datasets from five XFELs. Rugnux makes crystallographic data processing and structure solutions autonomous, fast, accessible, and cost-effective for both synchrotron and XFEL beamlines and individual researchers, at the rate and scale of modern MX experiments.

### Rugnux: from diffraction images to structure solution in one program

Rugnux is a new MX software package developed in the era of Artificial Intelligence (AI). It accelerates MX data processing and structure determination with GPUs, integrating the core computational steps and workflow orchestration typically distributed across multiple programs into a single codebase. Starting with raw diffraction images, it processes data, assigns symmetry, and assesses quality, then places the model, either an experimental structure^17^ or an AlphaFold prediction^18^,^19^, by molecular replacement and generates electron-density maps (Methods, Extended Data Fig. 1a). Rugnux processes rotation and still images with the same per-image 2D engine. For rotation data, the crystal is indexed once for the whole sweep, the geometry is post-refined against all images, and partial reflections are combined across the frames they span. For stills, each image is indexed and its orientation refined on its own, and partial intensities are scaled and merged across images (Extended Data Fig. 1b). Rugnux has no dependencies on external crystallographic software.

Rugnux performs per-frame analysis at kilohertz rates on a single consumer GPU and maintains the data in memory throughout the workflow, from the initial diffraction image to the resulting electron-density map. Its development employed large language models as tools for code exploration and implementation, with proposed changes evaluated against a curated validation suite of datasets. Operational robustness is achieved by validating automated decisions against the diffraction data throughout the processing and structure-determination workflow (Methods). All results presented here were obtained using default settings and the experimental parameters recorded in the image headers of the deposited datasets.

Rugnux includes a graphical user interface, the Jungfraujoch viewer, which displays diffraction images and runs complete data-processing jobs using the same computational engine and default settings as the command-line interface. Users can evaluate processing settings on an individual image before applying them to the entire dataset and export the corresponding command-line invocation for batch processing (Extended Data Fig. 2).

### One program, three operating modes

Rugnux is designed to deliver crystallographic results with low latency during data collection at beamlines, to enable fast data processing on a workstation, and to remain practical on conventional laboratory computers (Fig. 1). Its computationally intensive operations are accelerated on a GPU, while a CPU implementation allows the same program to run on hardware without a GPU (Extended Data Fig. 1).

**Figure 1.**
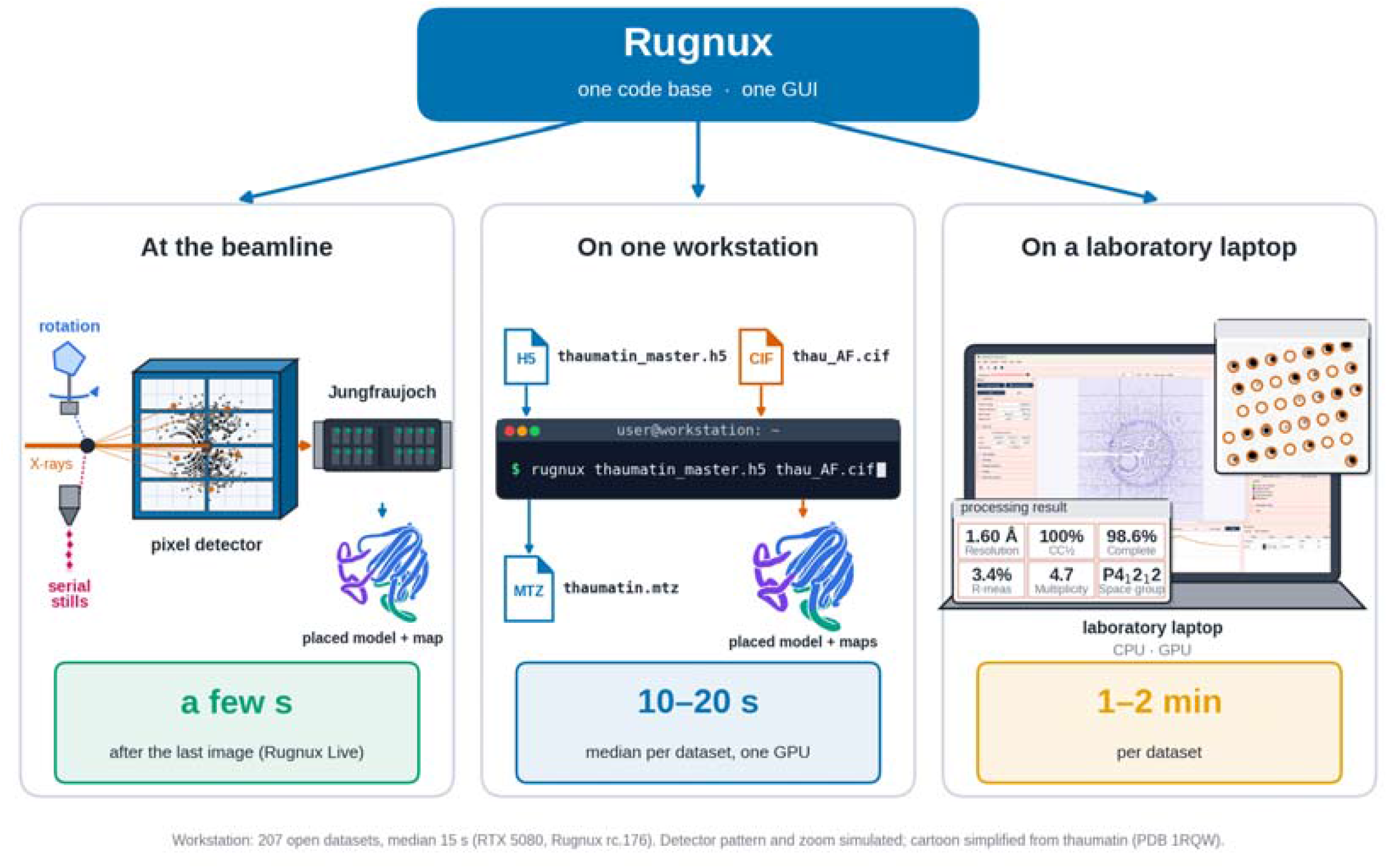
One codebase, zero dependencies, three operating modes. **Left**, at the beamline: diffraction images from a hybrid-pixel detector (PILATUS, EIGER) are streamed to a single Jungfraujoch server (FPGA + GPU), where Rugnux processes them as they arrive. **Center**, on one workstation: a single command takes the diffraction images and a model and produces merged data, the placed model, and electron density maps. **Right**, on a laboratory laptop: Rugnux viewer (GUI) drives the data processing and displays results.

#### At the beamline

We integrated Rugnux into the Jungfraujoch data acquisition system at MX beamlines at the upgraded Swiss Light Source 2.0. Frame-by-frame analysis starts as soon as data collection begins and keeps up with the detector data-rate. Final scaling/merging and MR start right after the shutter closes. We use a Jungfraujoch server (2x Intel Xeon 8558P; 4xNVIDIA L4) to transfer and perform on-the-fly analysis of diffraction data from an EIGER2 16M detector at PXII-X10SA. Using a cytochrome C test crystal, we collected a 360-degree dataset at 100 Hz frame-rate in 18 seconds, and obtained the final results within five seconds from the shutter closure, including processed data, the MR solution with AlphaFold models, and initial electron density maps (Extended Data Fig. 3). This short latency creates the possibility of using preliminary structural results to guide subsequent experimental decisions.

#### On one workstation

To assess processing performance and timing profiles across diverse datasets on a single workstation (AMD Ryzen 9 5950X, NVIDIA RTX 5080), we processed 207 publicly available datasets from 64 beamlines across 26 synchrotron facilities and 10 laboratory sources, using seven detector families (pan-facility test) (Fig. 2). Including disk reads and model-based validation (rigid body without MR), the median processing time was 15 s; 184 of 207 datasets completed within 30 s, and the slowest required 80 s (Fig. 2b). This outlier was a large unit-cell dataset to 1.39 Å resolution (PDB 9GDJ). Processing time increased with the total number of reflections, and the scaling step takes over as reflections grow (Fig. 2c, Extended Data Fig. 4). For the 201 macromolecular datasets with deposited structural models, subsequent rigid-body model-placement was completed within a few seconds on average.

**Figure 2.**
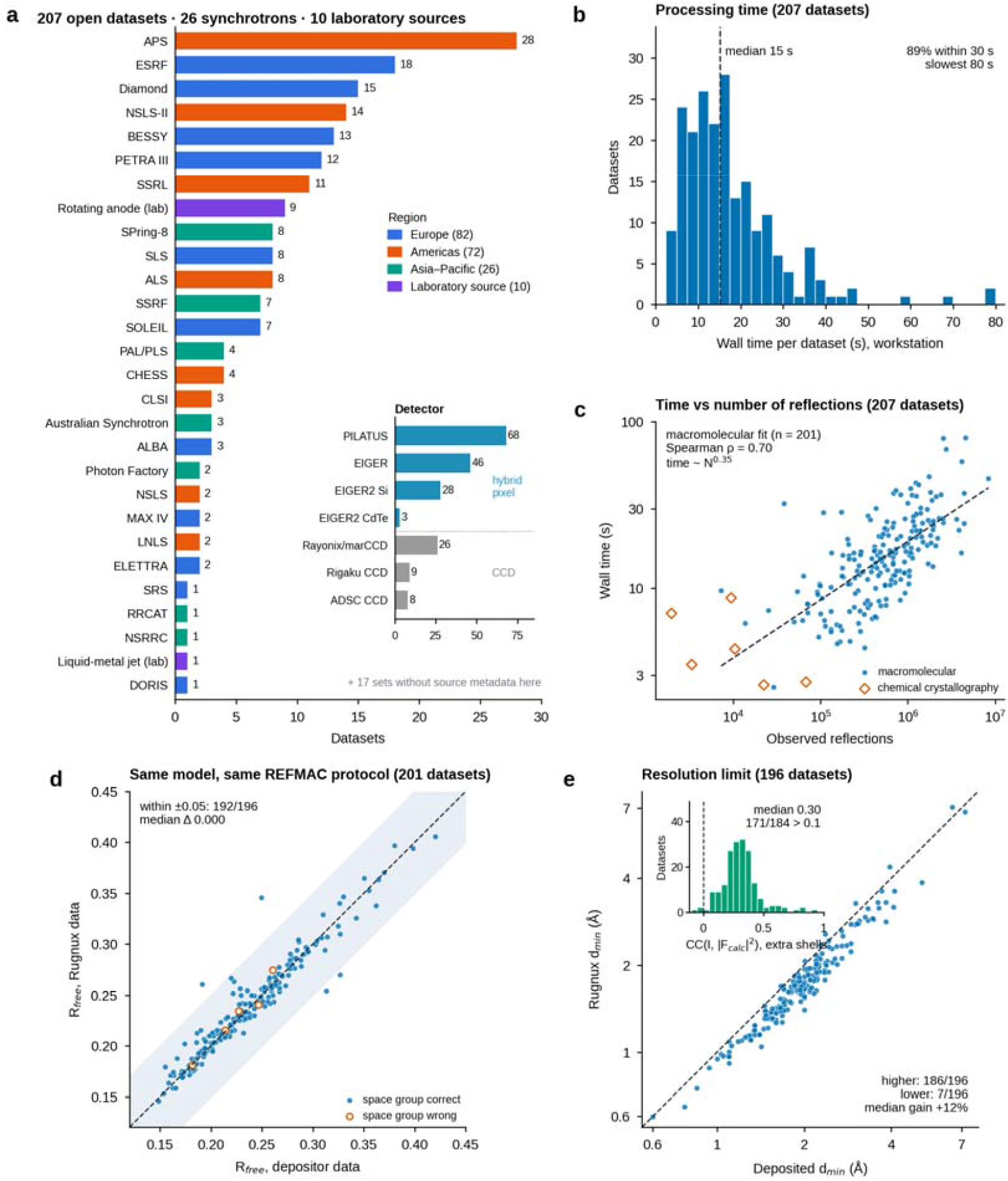
Pan-facility test at synchrotrons. **a**, The 20 open datasets (raw images of deposited PDB entries) per facility, colored by region; laboratory sources are rotating anodes and a liquid-metal jet. Inset, detector families. Eighteen sets without source metadata in the collection record are not shown. **b**, Data processing time dataset from files, including validation, on a desktop workstation (AMD Ryzen 9 5950X, NVIDIA GeForce RTX 5080). **c**, Data processing time against the number of observed reflections. **d**, R-free of each deposited model calculated by REFMAC with the same protocol against Rugnux data and against the depositor’s data, on the depositor’s free set; the dashed line marks identity and the shaded band ±0.05. **e**, High-resolution limit chosen by Rugnux against the deposited limit for 196 structures. Inset, histogram of the correlation between the intensities measured by Rugnux and |F_calc|^2^ from the deposited model, for reflections between the deposited and the Rugnux limits in 184 datasets where Rugnux produced higher resolution.

#### On a laboratory laptop

We next assessed how performance scaled across computing platforms. We processed four datasets from the pan-facility test (9ZLO, 9S02, 9GJX, and 4NWV), each ending with a structure placed by molecular replacement, on 11 configurations (Extended Data Table 1). The wall processing time varies from 10 sec to a few minutes. On a Windows laptop with GPU (Intel Core Ultra 7 255H; NVIDIA RTX PRO 2000), the total times are 38.6, 58.1, 47.9, and 156 sec, respectively.

Together, these results show that Rugnux combines low-latency processing at synchrotron data-collection rates with portability across computing platforms, letting users obtain crystallographic results during beamtime and revisit them in their own laboratory.

### Crystallographic data quality matches established programs across synchrotron facilities

We evaluated Rugnux against reference processing results using 201 macromolecular rotation datasets associated with deposited Protein Data Bank (PDB)^17^ structures. The datasets were the same pan-facility test set without chemical crystallography data (Fig. 2a). They spanned resolutions of 0.6–7.0 Å, unit-cell volumes of 2 × 10^4^ to 6 × 10^7^ Å^3^, and all seven crystal systems. For each dataset, we processed the raw diffraction images with Rugnux. 196 resulted in the correct space group. The five cases that ended with the wrong space group were due to twinning or pseudo-symmetry, and we reassigned their space groups to the deposited ones for refinement. We then refined the deposited structural model with REFMAC^20^ against both the resulting intensities and the reference data from the original deposition. We also used this dataset during Rugnux development, so it provides a broad assessment of performance across diverse experimental conditions rather than a fully independent validation set.

Rugnux-processed data reproduced the reference model quality in most cases. Among the 201 structures with an R-free comparison, 192 of 196 with the correct space group yielded R-free values within 0.05 of those calculated against the reference data. When space groups were corrected for the five Rugnux assigned different space groups, they also yield comparable R-free. The median ratio of R-free values from Rugnux and reference data was 1.00 (Fig. 2d). These results show that Rugnux produces data suitable for crystallographic refinement across a broad range of sources and experimental conditions, while symmetry assignment in pathological cases remains one limitation.

Rugnux also extended processing resolution beyond the deposited limit for 186 of 196 datasets, suggesting a conservative resolution cut-off was used in MX previously. Rugnux’s median limiting resolution was 1.69 Å, compared with 1.95 Å for the deposited data (Fig. 2e). For reflections beyond the deposited resolution limit (“extra shells”), the median correlation between the Rugnux observed intensities and structure factors calculated from the deposited model was 0.3 (Fig. 2e, inset), indicating that at least some of the additional high-resolution reflections contain useful signals.

### Fast and accurate processing of SFX data across XFEL facilities

Rugnux processes still images, as collected in serial crystallography, with the same per-image engine. Instead of following the rotation through the sweep, it indexes and integrates each image independently and merges partial intensities across images. We processed 11 public datasets from the Coherent X-ray Imaging Data Bank^21^, collected at five X-ray free-electron lasers (LCLS, SACLA, PAL-XFEL, EuXFEL, and SwissFEL) on five detector types (Rayonix MX340, MPCCD 4M, Rayonix MX225, AGIPD 1M, and JUNGFRAU 16M/4M; Extended Data Table 2). For SwissFEL data, we converted data to integer photon count format to allow compression^22^; otherwise, processing speed would be limited solely by disk reading time.

On the same workstation used in the synchrotron pan-facility test, from raw frames to merged data, a placed model and maps, Rugnux processed 140–674 frames s^−1^ at four of the facilities. That is 4.7 to 25 times faster than the frames were collected and above each facility’s operating repetition rate (Fig. 3, Extended Data Table 2). For European XFEL, with its higher repetition rate, the data processing rate reached 408–798 frames s^−1^, 1.4 to 5.3 times the rate at which these datasets were recorded. By optimizing the XFEL data format and compression and deploying higher-end GPUs, kilohertz processing rates are within reach. The single-server solution can also scale to match the higher repetition rates of next-generation XFELs.

**Figure 3.**
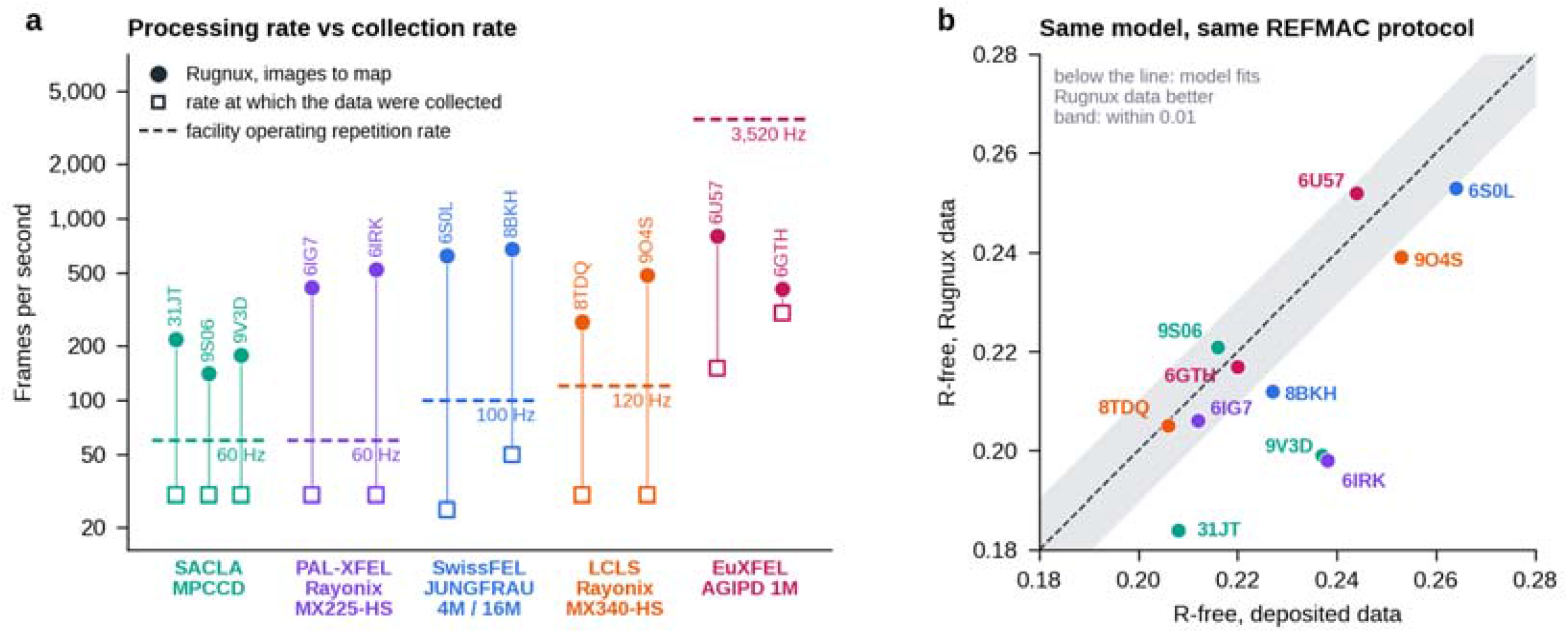
Pan-facility test at XFELs. Serial femtosecond crystallography datasets from five X-ray free-electron lasers processed by Rugnux on one workstation. **a**, Processing rate for each dataset, from raw frames to merged data, placed model and map coefficients (dots, labeled with the PDB entry), against the rate at which the dataset was collected (squares) and the facility’s operating repetition rate (dashed lines). The detector of each facility is given below its name. **b**, R-free of the deposited model, calculated against Rugnux data and against the deposited structure factors. Dashed line, identity; shaded band, ±0.01.

When calculated against the same deposited models using the same protocol, Rugnux data gave R-free values comparable to those of the deposited structure factors. The resolution limit was higher than the deposited for 9 of the 11. All together, Rugnux will make real-time data processing and structure determination a reality at current and future XFELs.

## Discussion

High-throughput crystallographic data processing has traditionally relied on computing clusters and pipelines that coordinate multiple software packages. Rugnux demonstrates an alternative: a single GPU-equipped workstation can process rotation datasets in seconds while retaining crystallographic data quality across a broad range of experimental sources, for both rotation and serial still data. Rugnux reduces dependence on distributed computing infrastructure and simplifies deployment at synchrotron beamlines. Rugnux has been deployed as a data-processing pipeline at MX beamlines at SLS 2.0 since August 2026, demonstrating its applicability to routine beamline operation.

Reducing processing latency also creates opportunities to change how crystallographic experiments are conducted. When data-quality estimates and initial structural information become available while a sample remains on the goniometer, subsequent data collection can be guided by the observations already obtained. Depending on the experiment, this could inform decisions about collecting additional images, measuring another crystal, or changing acquisition conditions. Such feedback moves toward adaptive experiments and, ultimately, more autonomous beamlines. After beamtime, researchers can also reprocess data on a laboratory workstation or laptop, giving them greater independence from facility computing resources and specialist support.

Rotation and serial still crystallography have long relied on separate software. The two methods differ mainly in what a single image records: a slice of a rotation, or a still that captures only part of each reflection. They otherwise share most of the processing chain. Because Rugnux handles both kinds of image with one per-image engine and one set of merging and validation steps, the same program and the same diagnostics can follow an experiment as it moves between the two, allowing systematic study of both experimental modes. Making both kinds of data quick to process on a single computer may lower the barrier to experiments that combine them, and let more groups treat serial and rotation data as complementary views of the same structure rather than as separate specialties. The current program already performs GPU-accelerated molecular replacement and rigid-body refinement. We are developing further GPU-based refinement capabilities, with the goal of producing refined structural models from diffraction images on the fly.

We envision a future in which the beamline evolves from a facility for collecting diffraction data into an environment for real-time structural discovery. Immediate structural feedback during data collection would allow researchers to steer measurements, test emerging hypotheses, and adapt experimental strategies as experiments unfold. We hope these capabilities will also open new avenues for exploring macromolecular dynamics, from probing ligand binding and temperature-dependent structural changes to capturing time-resolved processes, advancing both fundamental research and drug discovery.

Realizing this vision will require not only faster algorithms, but also software that makes the most of available computing resources. Facilities worldwide face rapidly rising IT costs, threatening their ability to deliver the full potential of upgraded beamlines in user experiments. We hope Rugnux, building on the work of many before us and bringing decades of crystallographic knowledge into a modern, highly efficient software framework, will help facilities deliver the full performance of their beamlines to the scientific community.

## Methods

### A. The program design

#### Overview

Rugnux takes diffraction images to merged intensities, a placed model and electron-density maps. A per-image loop runs on the GPU, and whole-dataset steps work on reflection lists (Extended Data Fig. 1). Rugnux builds on established methods, including, but not limited to, those of XDS^2^, MOSFLM^4^, DIALS^5^, CrystFEL^6^, POINTLESS and AIMLESS^23^,^24^, autoPROC^12^ and Phaser^25^, adapted and extended during development against experimental data. Ceres, FFTW^26^, cuFFT and GEMMI^27^ provide refinement, Fourier transforms, crystallographic symmetry operations and file handling, and per-step credits ship with the source code. The result is a standalone program for rotation and serial still crystallography that runs on a single computer.

#### Per-image processing

A pre-scan measures the beam center, beam-stop shadow, hot pixels, ice and powder rings, spot width and energy bandwidth. The beam center is found from the centrosymmetry of the mean image, computed by one FFT-based self-convolution at half-pixel spacing, which catches header errors of tens to hundreds of pixels. Each frame is then decoded on the GPU (bitshuffle-LZ4, CBF byte-offset and marCCD), azimuthally integrated, searched for spots, refined, predicted and integrated, and only a profile, a spot list and the integrated reflections return to the host. Integration is two-dimensional: each reflection is fitted on the frame where it is observed, with a profile learned from that frame’s strong spots, and partials are combined into full reflections during scaling. Because no frame waits for its neighbors, the loop can follow a live stream and is limited by the detector rather than by data transfer.

#### Indexing, refinement and passes

A lattice is accepted only if more spots fall on it than on the same lattice with the frames moved to another spindle angle. Distance, beam center, cell and orientation are refined on one half of the pass-1 reflections, and each step is kept only if the residual on the other half falls. A run is a tree of passes. Pass 1 works on the first half of the data. Pass 2 re-indexes and re-integrates the whole sweep at the refined geometry, and is repeated while it improves. Questions pass 1 cannot settle (beam center, lattice class, a doubled cell, a twin) run as alternative passes, and the better merge decides.

#### Scaling, symmetry and quality tests

Per-frame scales are smoothed with a cross-validated penalty. Absorption, detector and decay terms are kept only if they improve agreement on held-out data, and frames are rejected by ΔCC½^28^. Symmetry is scored operator by operator on normalized intensities (E^2^), and screw axes from systematic absences, zone by zone. Twinning is tested with the L test^29^ and translational pseudo-symmetry against shell-shuffled intensities^30^. The resolution limit is where CC½^31^ falls to 0.30 in a logistic fit. Where the data cannot decide (enantiomorphs, a screw axis on an unrecorded row, an inversion center hidden by Friedel’s law), Rugnux reports all options.

#### Molecular replacement and model validation

Rugnux places a model by molecular replacement. Fast rotation and translation searches run on the GPU, and the candidate placements are ranked by maximum likelihood. AlphaFold predicted models are prepared for the search: low-confidence residues are removed, B-factors are derived from pLDDT, and the model is split into rigid domains using the predicted aligned error^32^. The best placement is refined as a rigid body from low to high resolution, and a move is kept only if R-free improves. A placement is accepted only if it fits the data significantly better than the same model in random orientations. Only an accepted model may influence processing, by settling the hand, the indexing or the space group. Rugnux reports R-work, R-free and σA-weighted electron-density maps, plus an anomalous difference map where Bijvoet pairs were kept. Structure factors and bulk solvent are computed on GPU. Coordinates and B-factors are not refined.

#### Output and report

Rugnux writes merged and unmerged MTZ, mmCIF and SHELX HKLF4 files, and a report with a verdict from a closed vocabulary (OK, warnings, unusable, failed). The report also lists named pathology codes (for example ice rings, multiple lattices, translational pseudo-symmetry, anisotropy and twinning) and the evidence for each decision.

### B. Software

#### Implementation

Rugnux is written in C++ and CUDA and runs on Linux, Windows, and macOS. Every GPU step has a CPU counterpart, so a build without CUDA runs the whole pipeline, and on Apple silicon Bragg integration and part of molecular replacement run on Metal. Reductions that feed a decision are order-independent, so verdicts and space groups do not depend on thread count, number of GPUs, or compiler flags. Rugnux reads NXmx HDF5 and miniCBF files, with limited support for marCCD and SMV, and can follow a collection on the acquisition server as it is written.

#### Rugnux GUI/viewer

The GUI (jfjoch_viewer; Extended Data Fig. 2) is a Qt 6 application that links the same library. It draws images with spot, prediction, and resolution-ring overlays (Qt Graphics View) and plots per-image and merging statistics (Qt Charts). Processing jobs started from its settings panel run the Rugnux engine in a background thread and, with default settings, give the same result as the command line. Releases bundle the Qt runtime, so neither Qt nor a CUDA toolkit has to be installed.

#### Development with large language models

Rugnux was developed with large language models (Claude Code with Claude Opus 4.8, 5.0 and 5.5; Anthropic). We posed each problem, often a dataset that failed to process, and set the constraints; the model studied the data and the literature, implemented candidate solutions, and tested them on the affected datasets and then on the whole battery, against a baseline and independent references. We accepted a change only if it fixed the targeted cases without regressions elsewhere; about half of the ideas we tried were refuted by the data.

### C. Evaluation

#### Pan-facility test at synchrotrons

Development and benchmarking used a scripted battery. It holds 207 rotation datasets collected by other groups: 201 macromolecular datasets of deposited PDB entries and six chemical-crystallography datasets, from 26 synchrotrons and 10 laboratory sources. They come from eight public repositories: IRRMC (95 datasets)^33^, the SBGrid Data Bank (38)^34^, Zenodo (34)^35^, MXRDR (16)^36^, XRDa (5)^37^, the Keele University research data repository (4)^38^, the ESRF data portal (3)^39^ and UQ eSpace (1)^40^; the DOI of every dataset is listed with the Rugnux source code. They include cases of twinning, translational pseudo-symmetry and diffuse scattering, and controls for screw-axis assignment.

#### Pan-facility test at XFELs

We evaluated still-image processing on serial femtosecond crystallography data from five X-ray free-electron lasers (Extended Data Table 2). Other groups collected eleven datasets and deposited them in the Coherent X-ray Imaging Data Bank (CXIDB)^21^. The DOI of each dataset is listed with the Rugnux source code.

#### Comparison with deposited data

For the R-factor comparisons (Figs. 2d and 3b), each deposited model was scored with REFMAC^20^ 5.8 against Rugnux data and against the depositor’s structure factors with the same protocol. The Rugnux merge was reindexed into the model’s setting and reduced to the deposited space group, and both data sets were cut at the lower of the two resolution limits. Deposited intensities were converted to amplitudes by the French–Wilson method^41^ (ctruncate, CCP4^42^). R-free was read on the depositor’s free set at the first cycle of rigid-body refinement, before any shift, and entries that declare twinning were scored with twin refinement.

## Data availability

PDB identifiers, DOIs, and raw-data sources for all public benchmark datasets are listed with the source code, together with the battery manifests and the script that reproduces every comparison. All data processing results are available at https://doi.org/10.5281/zenodo.23596195.

## Code availability

Rugnux is available at https://gitea.psi.ch/mx/Jungfraujoch under GNU Public License. We provide binaries for Linux, Windows, and Mac OS.

## Acknowledgments

We thank the SLS macromolecular crystallography group for providing a stimulating environment for this work, in particular Vincent Olieric, Chia-Ying Huang, Guillaume Gotthard, Jiaxin Duan, Kiyofumi Takaba, and Oliver Bunk. We also thank SLS users for their valuable suggestions and for testing early versions of Rugnux. The coding, manuscript, and figure preparation were assisted by Claude from Anthropic Inc.

## Extended Data

**Extended Data Fig. 1.**
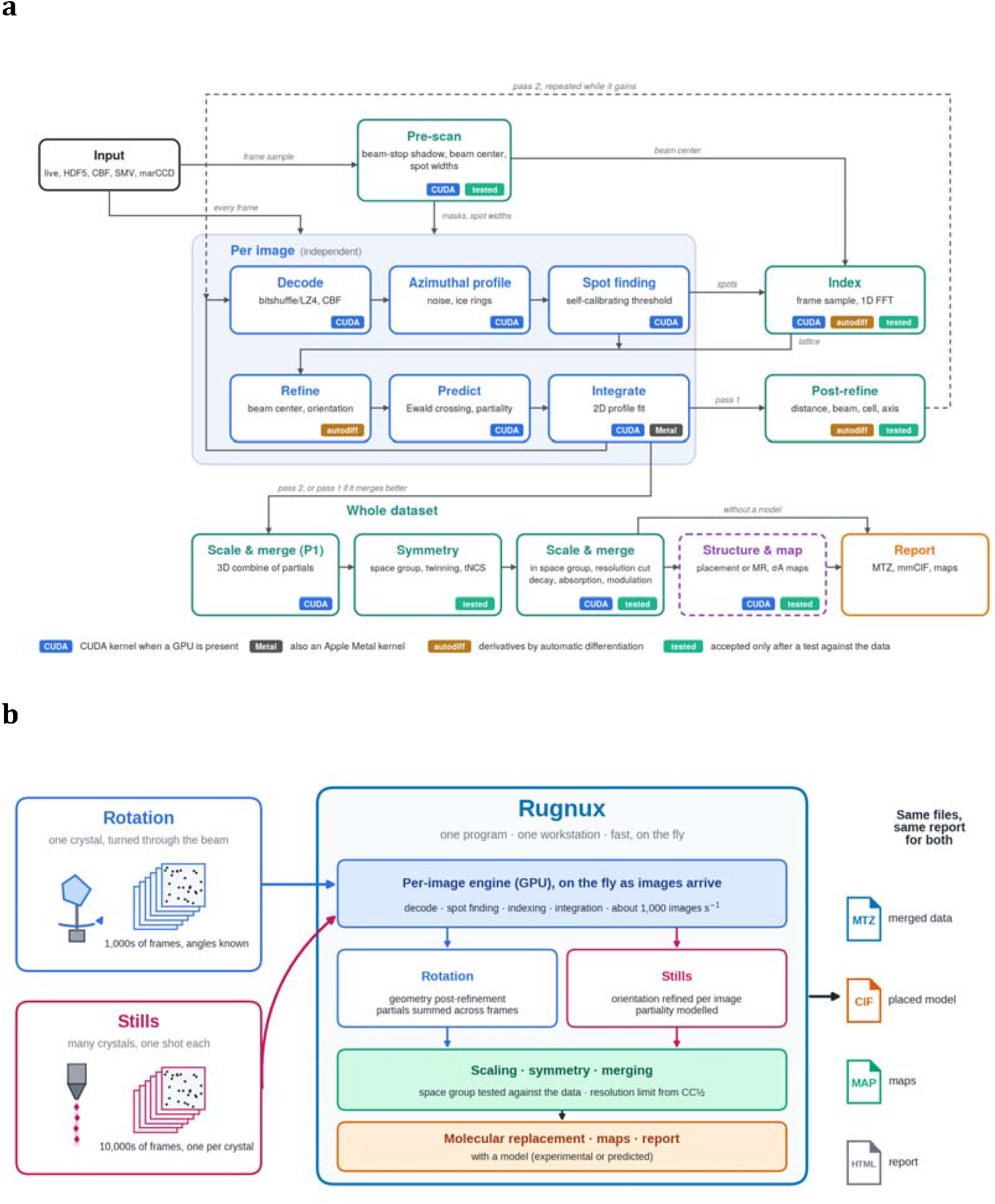
The Rugnux workflow. **a**, Rotation workflow. Each image is reduced to integrated reflections on the GPU as it arrives; indexing, geometry refinement and integration at the refined geometry follow, and merging, space-group assignment and model placement run on the reflection lists. The return arrow marks the refined geometry fed back for a second integration pass. **b**, Rotation and serial still images share the per-image engine and the merging and validation steps; rotation data are indexed once per sweep and post-refined, whereas stills are indexed and refined image by image.

**Extended Data Fig. 2.**
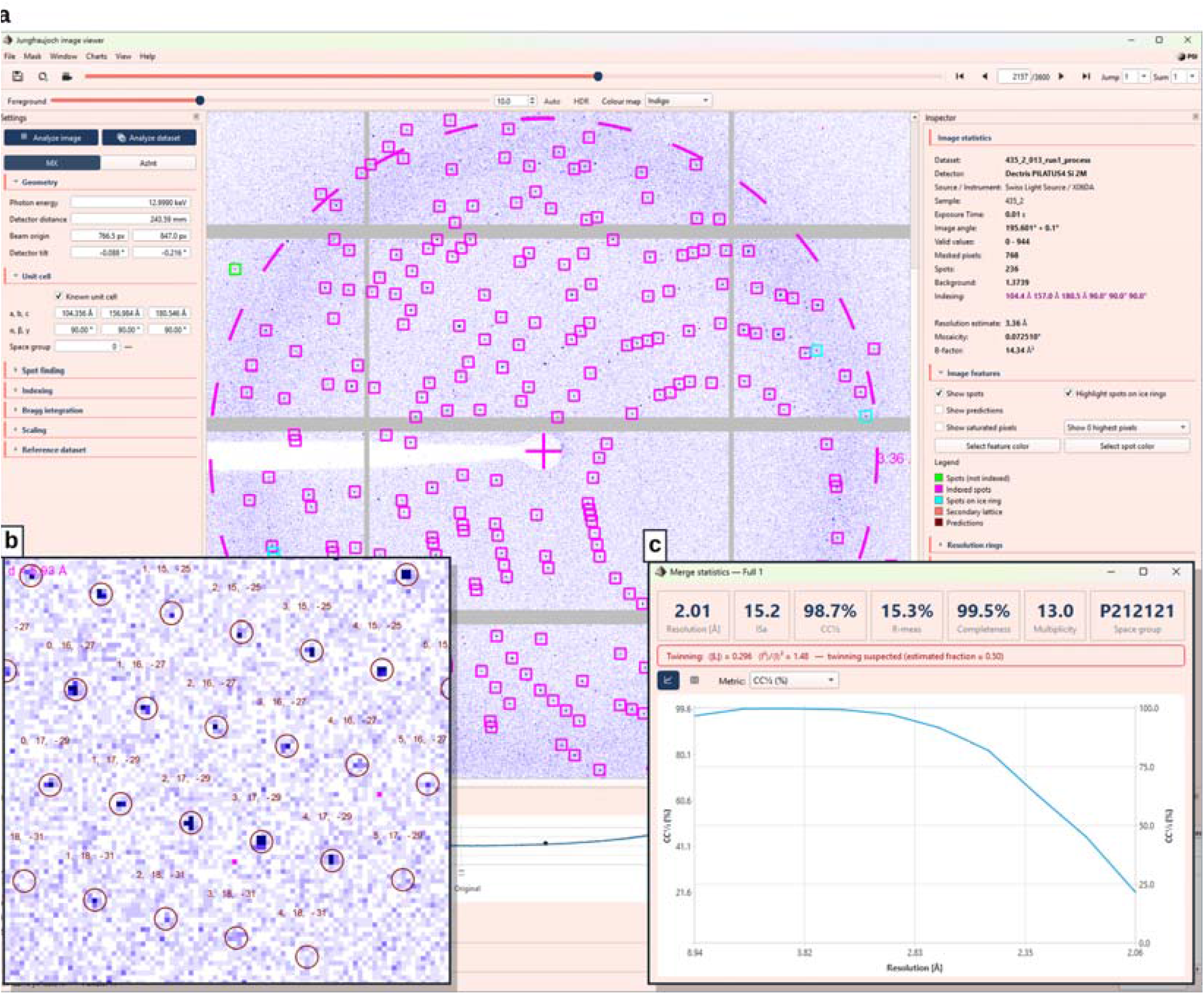
The Rugnux GUI (Jungfraujoch viewer). **a**, Main window for displaying and processing of a rotation dataset. Left, geometry and unit-cell settings; center, the diffraction image with spots found by Rugnux (magenta, indexed; green, not indexed; cyan, on an ice ring) and a resolution ring; right, per-image statistics. **b** (inset, lower left), zoomed view of the same dataset with predicted reflection positions (dark red circles) and their indices hkl. c (inset, lower right), merge statistics of the same dataset.

**Extended Data Fig. 3.**
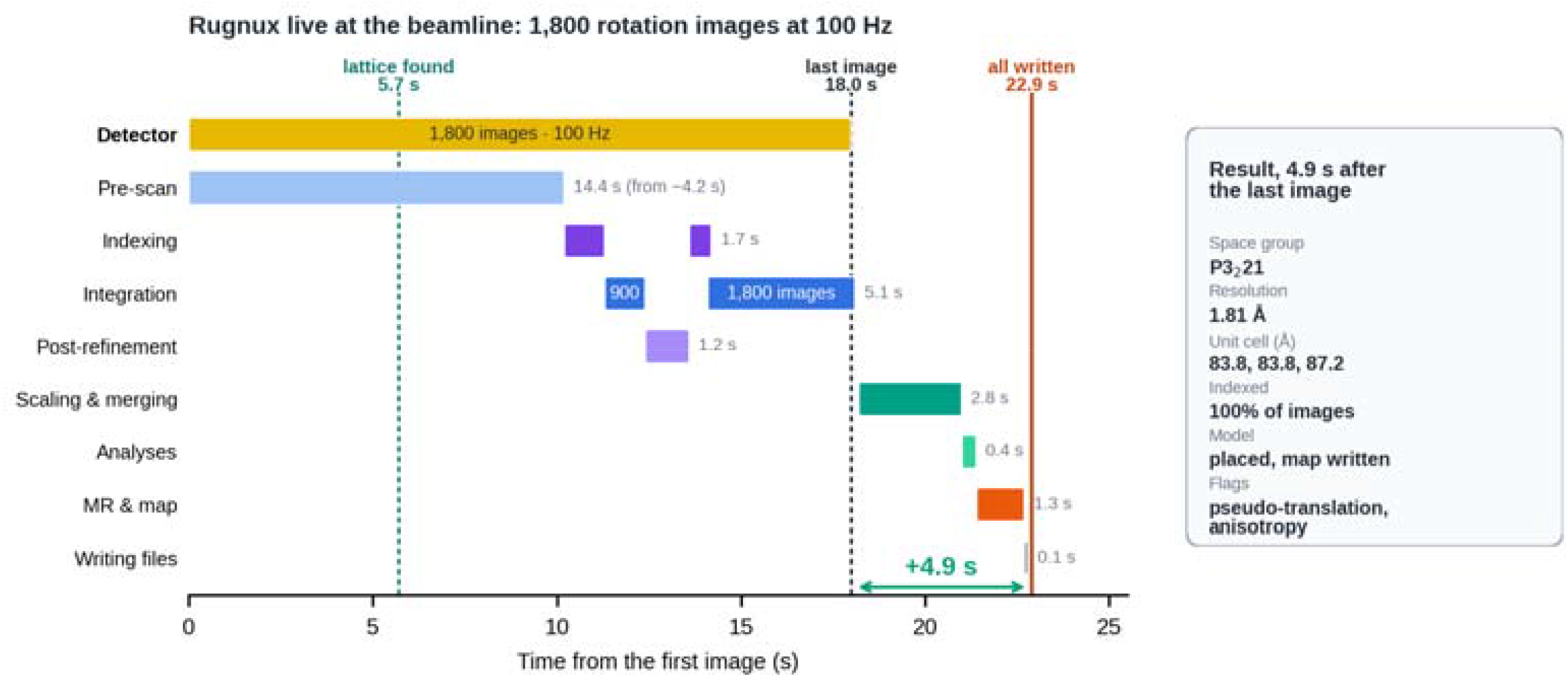
Rugnux Live performance at PXII-X10SA, SLS. Processing of a 1,800-image rotation dataset from a cytochrome C crystal during collection at beamline PXII-X10SA, SLS 2.0 (October 21, 2026), recorded at 100 Hz (0.2 deg per 0.01 sec) on an EIGER2 16M detector. A lattice was found from the first 60° at 5.7 s, and indexing, integration of the first 900 images, post-refinement and re-indexing at the refined geometry follow. Then integration of all 1,800 images at the refined geometry was catching up with data collection. After the last image, only scaling, symmetry determination and merging, data-quality analyses, molecular replacement with map calculation and file writing remain, which took 4.9 seconds.

**Extended Data Fig. 4.**
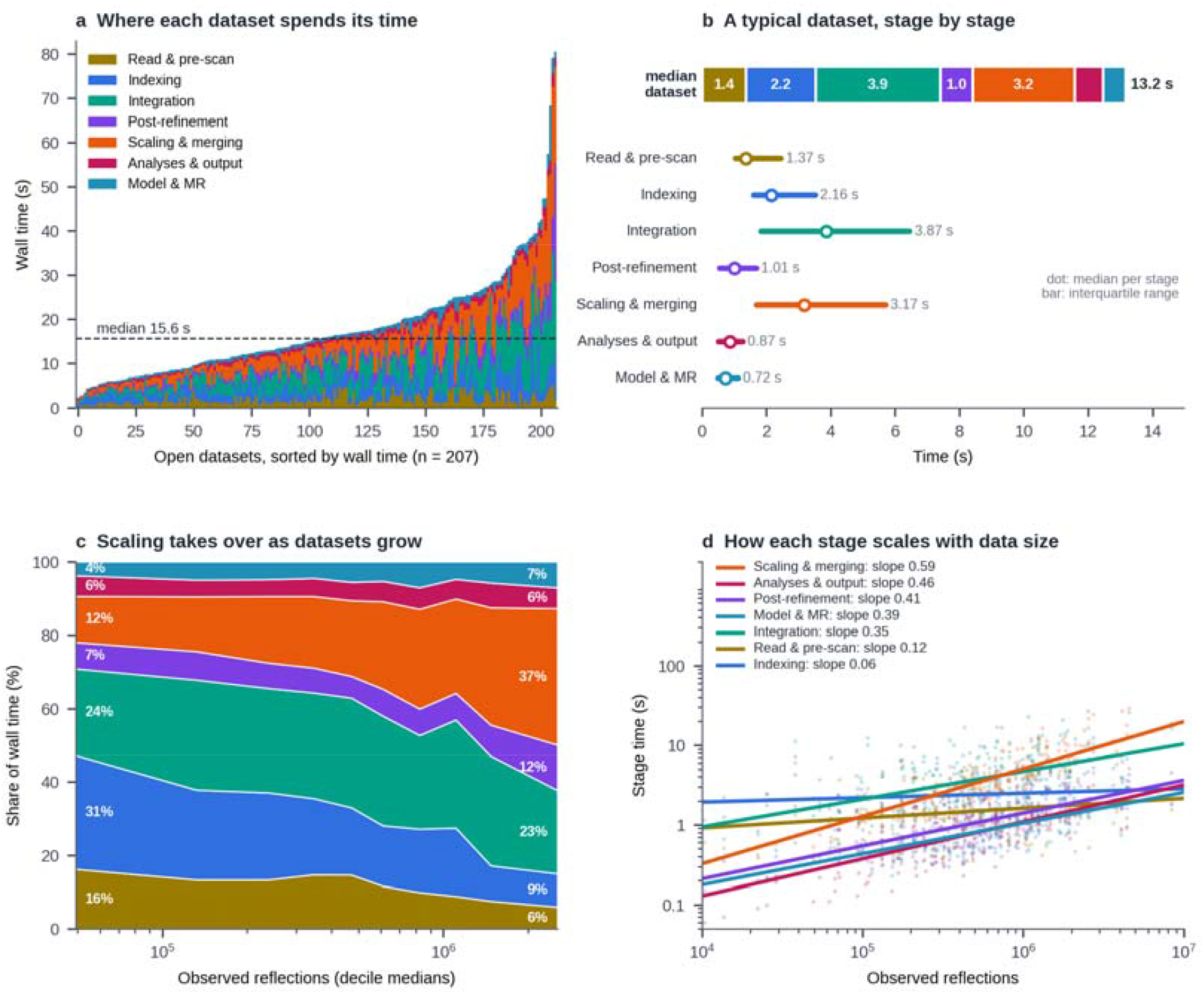
Timing statistics of 207 synchrotron pan-facility tests. All 207 datasets were processed with Rugnux on one workstation with an RTX 5080 GPU, with no other jobs on the GPU. Each dataset ran end to end with the deposited model supplied, where one existed. **a**, Processing time per dataset, split into processing stages. The dashed line marks the median. **b**, Stage-by-stage timeline of a median dataset. Dots mark the median time per stage and bars the interquartile range. **c**, Each stage’s share of the total processing time against the number of observed reflections, using the median of each decile of datasets. **d**, Time per stage against the number of observed reflections (log–log). Lines show power-law fits.

**Extended Data Table 1.**
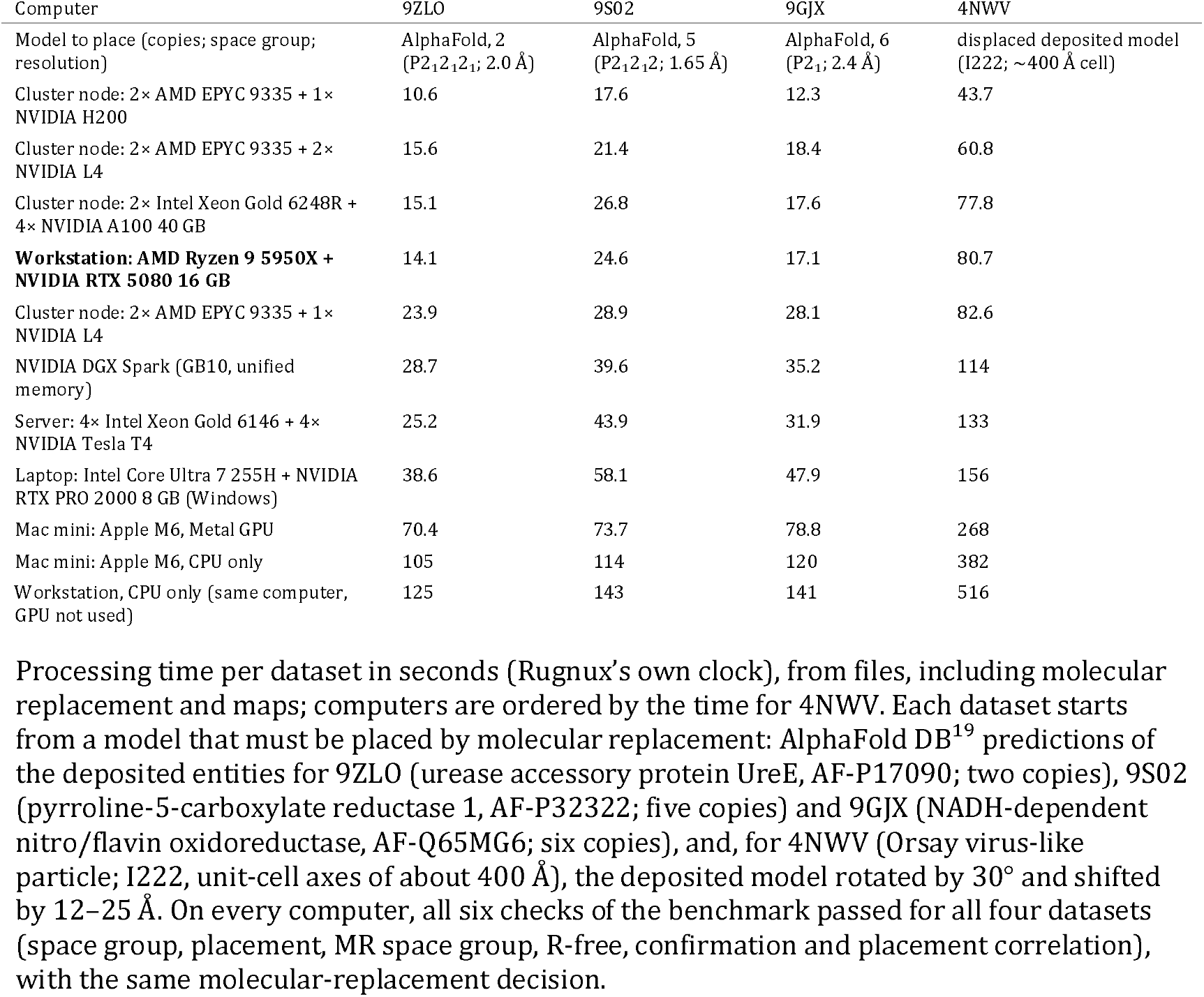
Processing time on different computers, from images to a structure placed by molecular replacement.

**Extended Data Table 2.**
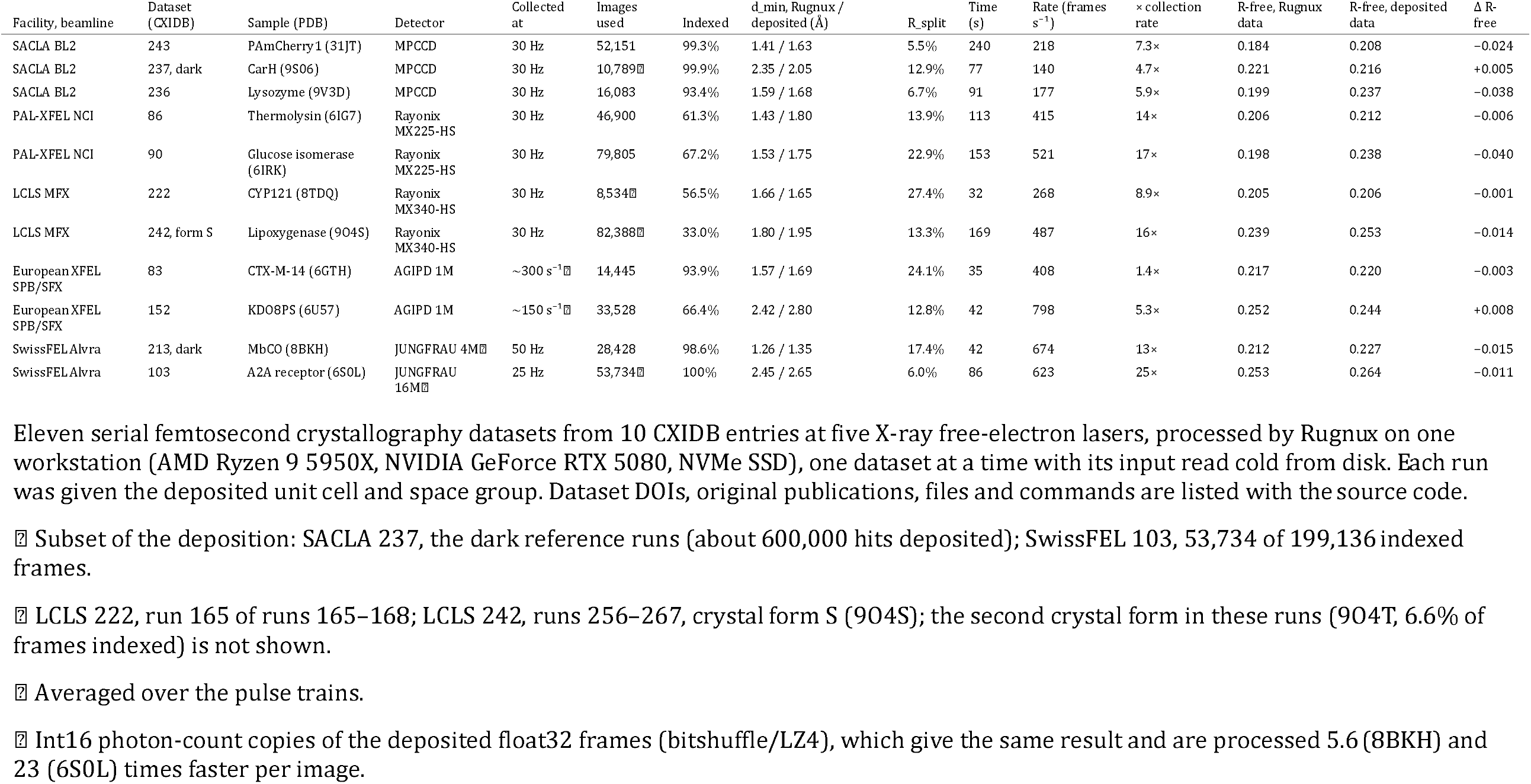
Serial (still-image) datasets from XFELs processed with Rugnux.

